# Exploring the genetic architecture of inflammatory bowel disease by whole genome sequencing identifies association at *ADCY7*

**DOI:** 10.1101/058347

**Authors:** Yang Luo, Katrina M. de Lange, Luke Jostins, Loukas Moutsianas, Joshua Randall, Nicholas A. Kennedy, Christopher A. Lamb, Shane McCarthy, Tariq Ahmad, Cathryn Edwards, Eva Goncalves Serra, Ailsa Hart, Chris Hawkey, John C. Mansfield, Craig Mowat, William G. Newman, Sam Nichols, Martin Pollard, Jack Satsangi, Alison Simmons, Mark Tremelling, Holm Uhlig, David C. Wilson, James C. Lee, Natalie J. Prescott, Charlie W. Lees, Christopher G. Mathew, Miles Parkes, Jeffrey C. Barrett, Carl A. Anderson

## Abstract

In order to further resolve the genetic architecture of the inflammatory bowel diseases, ulcerative colitis and Crohn’s disease, we sequenced the whole genomes of 4,280 patients at low coverage, and compared them to 3,652 previously sequenced population controls across 73.5 million variants. To increase power we imputed from these sequences into new and existing GWAS cohorts, and tested for association at ~12 million variants in a total of 16,432 cases and 18,843 controls. We discovered a 0.6% frequency missense variant in *ADCY7* that doubles risk of ulcerative colitis, and offers insight into a new aspect of disease biology. Despite good statistical power, we did not identify any other new low-frequency risk variants, and found that such variants as a class explained little heritability. We did detect a burden of very rare, damaging missense variants in known Crohn’s disease risk genes, suggesting that more comprehensive sequencing studies will continue to improve our understanding of the biology of complex diseases.

## Introduction

Crohn’s disease and ulcerative colitis’ the two common forms of inflammatory bowel disease (IBD), are chronic and debilitating diseases of the gastrointestinal tract that result from the interaction of environmental factors’ including the intestinal microbiota, with the host immune system in genetically susceptible individuals. Genome-wide association studies (GWAS) have identified 210 IBD associated loci that have substantially expanded our understanding of the biology underlying these diseases^1–7^. The correlation between nearby common variants in human populations underpins the success of the GWAS approach, but this also makes it difficult to infer precisely which variant is causal, the molecular consequence of that variant, and often even which gene is perturbed. Rare variants, which plausibly have larger effect sizes, can be more straightforward to interpret mechanistically because they are correlated with fewer nearby variants. However, it remains to be seen how much of the heritability^8^ of complex diseases is explained by rare variants. Well powered studies of rare variation in IBD thus offer an opportunity to better understand both the biological and genetic architecture of an exemplar complex disease.

The marked drop in the cost of DNA sequencing has enabled rare variants to be captured at scale, but there remains a fundamental design question regarding how to most effectively distribute short sequence reads in two dimensions: across the genome, and across individuals. The most important determinant of GWAS success has been the ability to analyze tens of thousands of individuals, and detecting rare variant associations will require even larger sample sizes^9^. Early IBD sequencing studies concentrated on the protein coding sequence in GWAS-implicated loci^10–13^, which can be naturally extended to the entire exome^14–16^. However, coding variation explains at most 20% of the common variant associations in IBD GWAS loci^17^, and others have more generally observed^18^ that the substantial majority of complex disease associated variants lie in non-coding, presumed regulatory, regions of the genome. Low coverage whole genome sequencing has been proposed^19^ as an alternative approach that captures this important noncoding variation, while being cheap enough to enable thousands of individuals to be sequenced. As expected, this approach has proven valuable in exploring rarer variants than those accessible in GWAS^20,21^, but is not ideally suited to the analysis of extremely rare variants.

Our aim was to determine whether low coverage whole genome sequencing provides an efficient means of interrogating these low frequency variants, and how much they contribute to IBD susceptibility. We present an analysis of the whole genome sequences of 4,280 IBD patients, and 3,652 population controls sequenced as part of the UK10K project^22^, both via direct comparison of sequenced individuals and as the basis for an imputation panel in an expanded UK IBD GWAS cohort. This study allows us to examine, on a genome-wide scale, the role of low-frequency (0.1%≥ MAF < 5%) and rare (MAF < 0.1%) variants in IBD risk (Figure 1).

**Figure 1.**
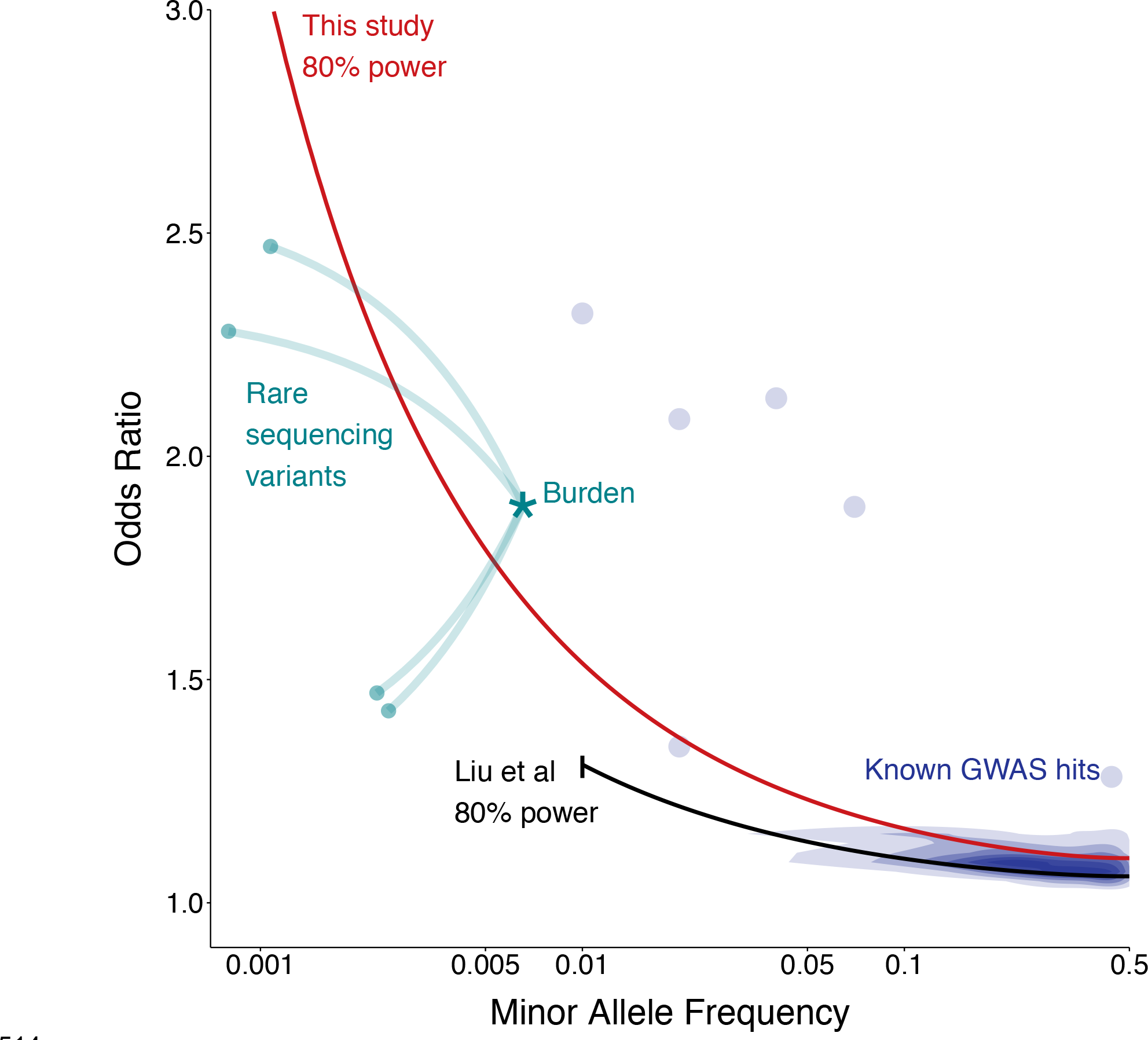
Relative power of this study compared to previous GWAS. Black line shows the path through frequency-odds ratio space where the latest IIBDGC meta-analysis had 80% power. Red line shows the same for this study. The earlier study had more samples but restricted their analysis to MAF > 1%. Purple density and points show known GWAS loci, green points show known ***NOD2*** rare variants, and the star shows their equivalent position when tested by gene burden, rather than individually.

## Results

### Whole genome sequencing of 8,354 individuals

We sequenced the whole genomes of 2,697 Crohn’s disease patients (median coverage 4x) and 1,817 ulcerative colitis patients (2x), and jointly analyzed them with 3,910 population controls (7x) sequenced as part of the UK10K project^22^ (Figure 2). We discovered 87 million autosomal single nucleotide variants (SNVs) and 7 million short indels (Supplementary Methods). We then applied support vector machines for SNVs and GATK VQSR^23^ for indels to distinguish true sites of genetic variation from sequencing artifacts (Figure 2, Supplementary Methods). We called genotypes jointly across all samples at the remaining sites, followed by genotype refinement using the BEAGLE imputation software^24^. This procedure leverages information across multiple individuals and uses the correlation between nearby variants to produce high quality data from relatively low sequencing depth. We noted that genotype refinement was locally affected by poor quality sites that failed further quality control analyses (Supplementary Methods), so we ran BEAGLE a second time after these exclusions, yielding a set of 76.7 million high quality sites. Over 99% of common SNVs (MAF ≥ 5%) were also found in 1000 Genomes Project Phase 3 Europeans, indicating high specificity. Among rarer variants, 54.6 million were not seen in 1000 Genomes, demonstrating the value of directly sequencing the IBD cases and UK population controls. Additional sample quality control (Supplementary Methods) left a final dataset of 4,280 IBD patients and 3,652 controls.

**Figure 2.**
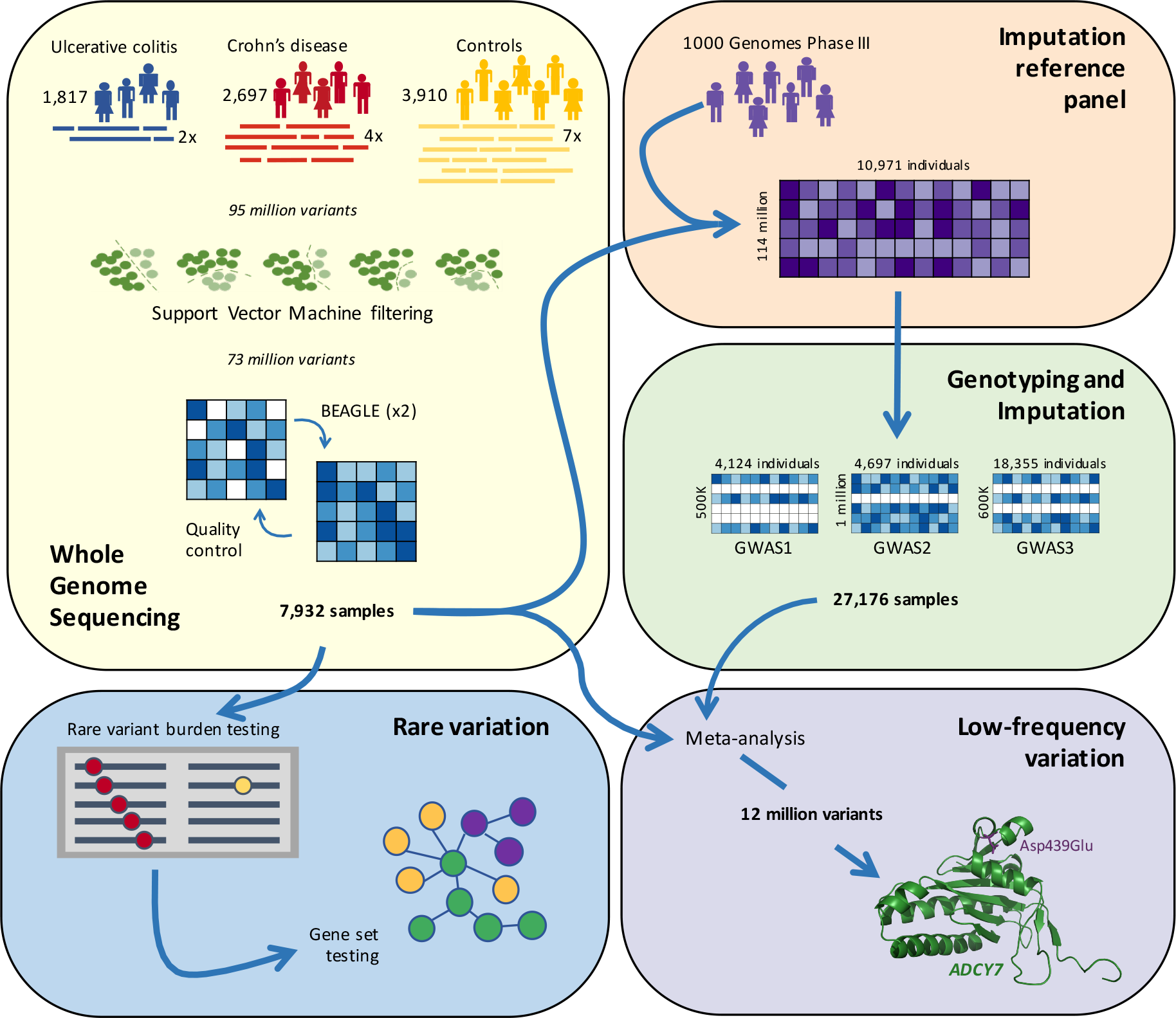
Overview of our study. Variants were called from raw sequence reads in three groups of samples, and jointly filtered using support vector machines. The resulting genotypes were refined using BEAGLE and incorporated into the reference panel for a GWAS-imputation based meta-analysis, which discovered a low frequency association in ADCY7. A separate gene-based analysis identified a burden of rare damaging variants in certain known Crohn’s disease genes. The partial predicted crystal structures for ADCY7 is obtained from the SWISS-MODEL respository.

Across these individuals we also discovered 180,000 deletions, duplications and multiallelic copy number variants (CNVs) using GenomeStrip 2.0^25^, but noted large differences in sensitivity between the three different sample sets (Supplementary Figure 5). Following quality control (Supplementary Methods), including removal of CNVs with length < 60 kilobases, we observed an approximately equal number of variants in cases and controls, but retained only 1,467 CNVs. Even after this stringent filtering, we still observed a genome-wide excess of rare CNVs in controls (P=0.006), suggesting that high coverage whole genome sequencing balanced in cases and controls will be required to evaluate the contribution of rare structural variation to IBD risk.

We individually tested 13 million SNVs, small indels and SVs with MAF ≥0.1% for association, and observed that we had successfully eliminated systematic differences due to sequence depth (λ_1000___UC_ = 1.05, λ_1000___CD_ = 1.04, λ_1000JBD_ =1.06, Supplementary Figure 6), while still retaining power to detect known associations. However, even this stringent quality control could not eliminate all false positives: we saw extremely significant p-values at many SNPs outside of known loci (e.g. ~7,000 with p < 10“^−15^), 95% of which had an allele frequency below 5% (Supplementary Figure 5). While we estimate that this analysis produced well calibrated association test statistics for more than 99% of sites, the heterogeneity of our sequencing depths makes it challenging to differentiate between false-positives driven by remaining artifacts and true low-frequency associations.

### Imputation into GWAS

Even had we been able to fully remove biases introduced by differences in sequencing depth, our WGS dataset alone would not have been well powered to identify associations missed by previous studies. We therefore built a phased reference panel of 10,971 individuals from our low coverage whole genome sequences and 1000 Genomes Phase 3 haplotypes (Supplementary Methods), in order to use imputation to leverage IBD GWAS to increase our power. Previous data have shown that such expanded reference panels significantly improve imputation accuracy of low-frequency variants^26^, and because our GWAS cases and controls were genotyped using the same arrays, they should be not be differentially affected by the variation in sequencing depths in the reference panel.

We next generated a new UK IBD GWAS dataset by genotyping 8,860 IBD patients without previous GWAS data and combining them with 9,495 UK controls from the Understanding Society project (www.understandingsociety.ac.uk), all genotyped using the Illumina HumanCoreExome v12 chip. We then added previous UK IBD GWAS samples that did not overlap with those in our sequencing dataset^27,28^. Finally, we imputed all of these samples using the PBWT^29^ software and the reference panel described above, and combined these imputed genomes with our sequenced genomes to create a final dataset of 16,432 IBD cases and 18,843 UK population controls. This imputation produced high quality genotypes at 12 million variants that passed typical GWAS quality control (Supplementary Methods), and represented more than 90% of sites with MAF >0.1% that we could directly test in our sequences. Compared to the most recent meta-analysis by the International IBD Genetics Consortium^30^, which used a reference panel almost ten times smaller than ours, we tested an additional 2.5 million variants for association to IBD.

### Asp439Glu in *ADCY7* doubles risk of ulcerative colitis

This analysis revealed four previously undescribed genome-wide significant IBD associations’ three of which had MAF > 10%, so we carried them forward to a meta-analysis of our data and published IBD GWAS summary statistics^31^. The fourth (P = 3x10“^12^) was a 0.6% missense variant (Asp439Glu, rs78534766) in ***ADCY7*** that doubles risk of ulcerative colitis (OR=2.24, 95% CI =1.79-2.82), and is strongly predicted to alter protein function (SIFT = 0, PolyPhen = 1, MutationTaster = 1). A previous report described an association between an intronic variant in this gene and Crohn’s disease^32^, but our signal at this variant (P = 2.9×10^−7^) vanishes after conditioning on the nearby associations at *NOD2*, (conditional P = 0.82). By contrast, we observed that Asp439Glu shows nominal association with Crohn’s disease after conditioning on ***NOD2*** (P = 7.5×10^−5^, 0R=1.40), while the significant signal remains for ulcerative colitis (Figure 3). Thus’ the strongest single alleles associated to both Crohn’s disease and ulcerative colitis (outside the major histocompatibility complex) affect genes that are, apparently coincidentally, only 30 kilobases apart (Figure 3).

**Figure 3.**
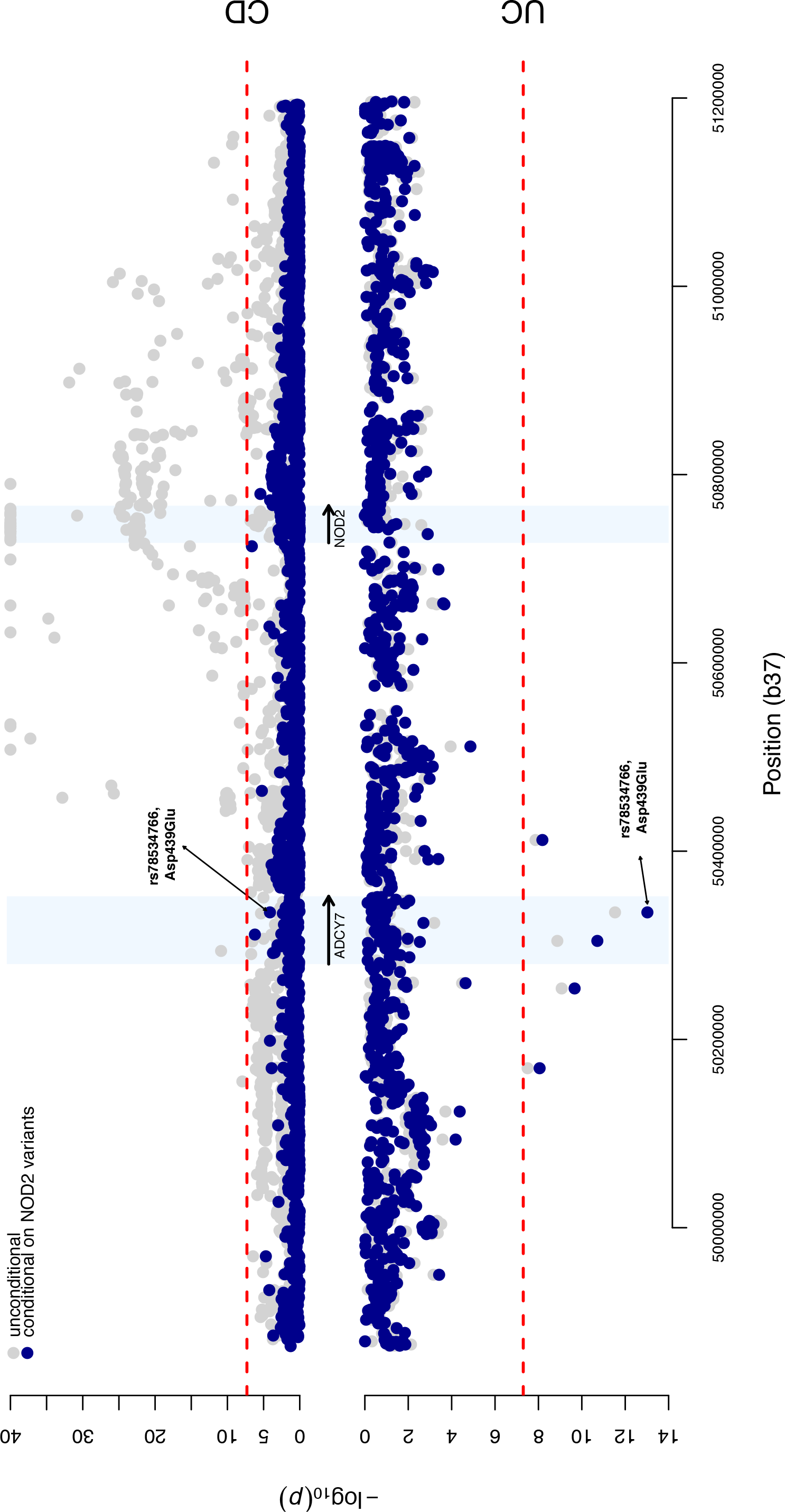
Association analysis for the *NOD2/ADCY7* region in chromosome 16. Results from the single variant association analysis are presented in gray, and conditioned on seven known ***NOD2*** risk variants in blue. Results for CD are shown on the top half, UC on the bottom half. Dashed red lines indicate genome-wide significance, at α=5e-08.

The protein encoded by *ADCY7*, adenylate cyclase 7, is one of a family of ten enzymes that convert ATP to the ubiquitous second messenger cAMP. Each has distinct tissue-specific expression patterns’ with ***ADCY7*** being expressed in haemopoietic cells. Here, cAMP modulates innate and adaptive immune functions’ including the inhibition of the pro-inflammatory cytokine TNFa, itself the target of the most potent current therapy in IBD^33^. Indeed, myeloid-specific Adcy7 knockout mice (constitutive knockouts die in utero) show higher stimulus-induced production of TNFa by macrophages’ impairment in B cell function and T cell memory, an increased susceptibility to LPS-induced endotoxic shock, and a prolonged inflammatory response^34,35^. In human THP-1 (monocyte-like) cells’ siRNA knockdown of ***ADCY7*** also leads to increased TNFa production.^36^. Asp439Glu affects a highly conserved amino acid in a long cytoplasmic domain immediately downstream of the first of two active sites and may affect the assembly of the active enzyme through misalignment of the active sites^37^.

### Low-frequency variation makes a minimal contribution to IBD susceptibility

The associated variant in ***ADCY7*** represents precisely the class of variant (below 1% MAF, OR ~2) that our study design was intended to probe, making it notable as our single discovery of this type. We had 66% power to detect that association, and reasonable power even for more difficult scenarios (e.g. 29% for 0.2% MAF and OR=2, or 11 % for 0.5% MAF and OR=1.5). As noted by others^38^, heritability estimates for low frequency variants as a class are exquisitely sensitive to potential bias from technical and population differences. We therefore analyzed only the imputed GWAS samples to eliminate the effect of differential sequencing depth, and applied a more stringent SNP and sample quality control (Supplementary Methods). We used the restricted maximum likelihood (REML) method implemented in GCTA^39^ and estimated that autosomal SNPs with MAF > 0.1% explain 28.4% (s.e. 0.016) and 21.1% (s.e. 0. 012) of the variation in liability for Crohn’s and ulcerative colitis, respectively. Despite SNPs with MAF < 1% representing approximately 81% of the variants included in this analysis, they explained just 1.5% of the variation in liability. While these results are underestimates due to limitations of our data and the REML approach, it seems very unlikely that a large fraction of IBD risk is captured by variants like ***ADCY7*** Asp439Glu. Thus, our discovery of ***ADCY7*** actually serves as an illustrative exception to a series of broader observations^40^ that low-frequency, high-risk variants are unlikely to be important contributors to IBD risk.

### The role of rare variation in IBD risk

Our low coverage sequencing approach does not perfectly capture very rare and private variants because the cross-sample genotype refinement adds little information at sites where nearly all individuals are homozygous for the major allele. Similarly, these variants are difficult to impute from GWAS data: even using a panel of more than 32,000 individuals offers little imputation accuracy below 0.1% MAF^26^. Thus, while our sequence dataset was not designed to study rare variants, it is the largest to date in IBD, and has sufficient specificity and sensitivity to warrant further investigation (Supplementary Figure 7). Because enormous sample sizes would be required to implicate any single variant, we used a standard approach from exome sequencing^41^, where variants of a particular functional class are aggregated into a gene-level test. We extended Derkach *et al*’s Robust Variance Score statistic^42^ to account for our sequencing depth heterogeneity, because existing rare variant burden methods gave systematically inflated test statistics.

For each of 18,670 genes, we tested for a differential burden of rare (MAF < 0.5%, excluding singletons) functional or predicted damaging coding variation in our sequenced cases and controls (Online Methods, Supplementary Table 6). We detected a significant burden of damaging rare variants in the well-known Crohn’s disease risk gene ***NOD2*** (P_functional_ = 1×10^−7^, Supplementary Figure 9), which was independent of the known low-frequency ***NOD2*** risk variants (Online Methods). We noted that the additional variants (Figure 4) that contribute to this signal explain only 0.13% of the variance in disease liability, compared to 1. 15% for the previously known variants^10^, underscoring the fact that very rare variants cannot account for much population variability in risk.

**Figure 4.**
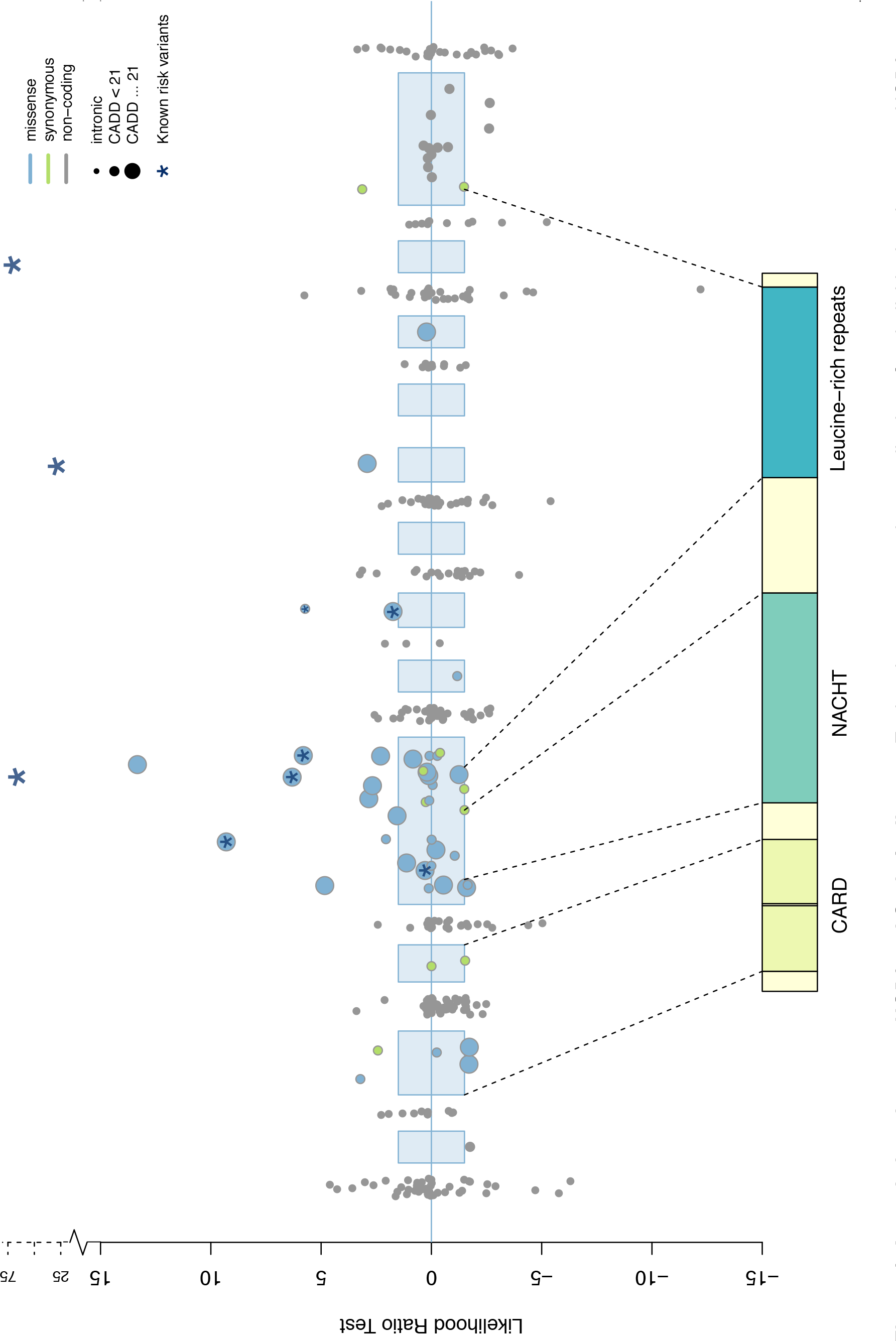
Associations between *N0D2* and Crohn’s disease. Each point represents the contribution of an individual variant to our ***N0D2*** burden test. Three common variants (rs2066844, rs2066845, rs2066847) are shown for scale, and the six rare variants identified by targeted sequencing are starred. Exonic regions (not to scale) are marked in blue, with their corresponding protein domains highlighted.

Some genes implicated by IBD GWAS had suggestive p-values, but did not reach exome-wide significance (P=5×10^−7^), so we combined individual gene results into two sets: (i) 20 genes that had been confidently implicated in IBD risk by fine-mapping or functional data, and (ii) 63 additional genes highlighted by less precise GWAS annotations (Supplementary Methods, Supplementary Table 9). We tested these two sets (after excluding ***NOD2***, which otherwise dominates the test) using an enrichment procedure^41^ that allows for differing direction of effect between the constituent genes (Supplementary Methods). We found a burden in the twelve confidently implicated Crohn’s disease genes that contained at least one damaging missense variant (Pd_amaging_ = 0.0045). By contrast, we saw no signal in the second, more generic set of genes (P=0.94, Figure 5, Table 1).

**Figure 5.**
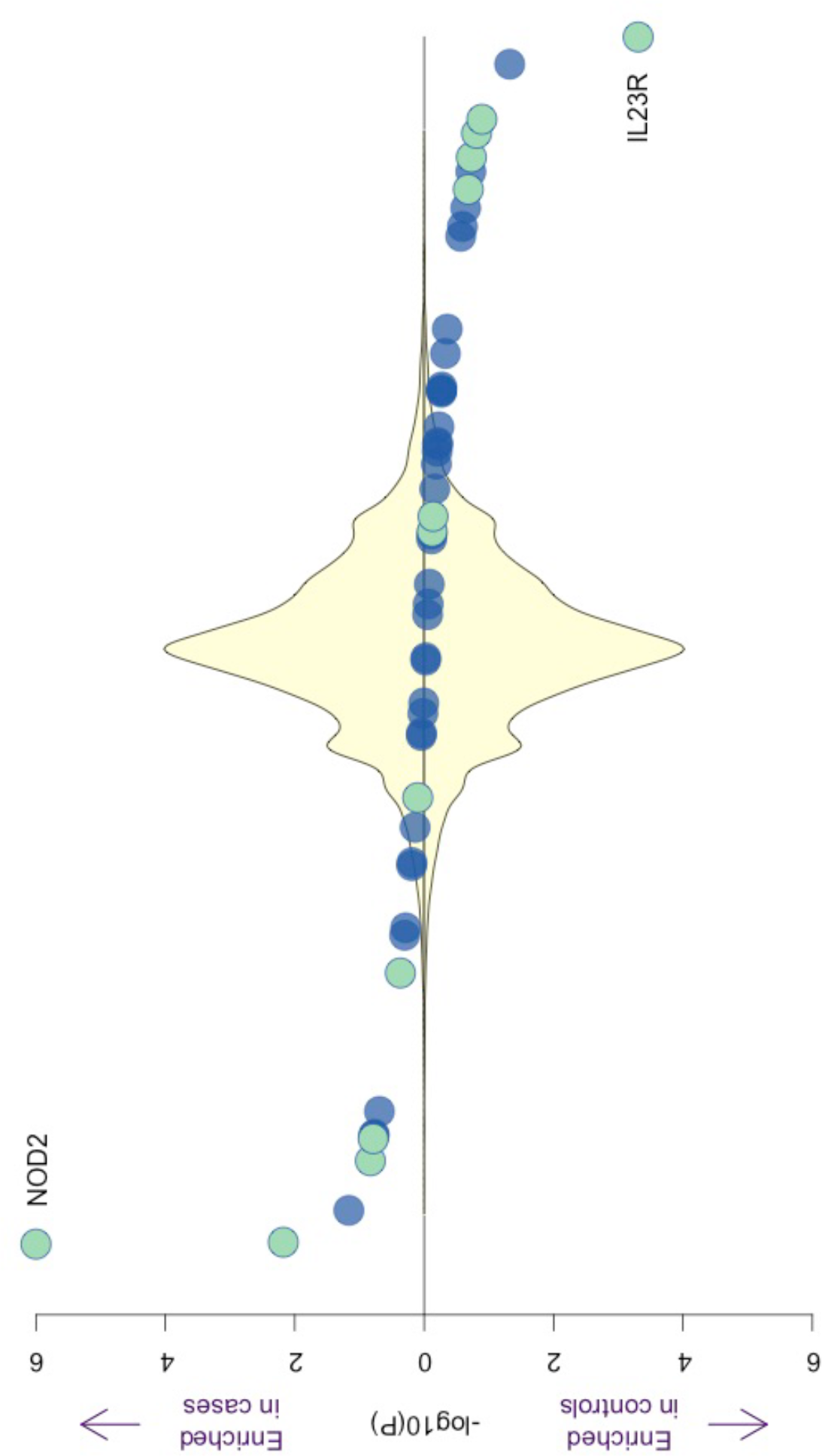
Burden of rare damaging variants in Crohn’s disease. Each point represents a gene in our confidently implicated (green) or generically implicated (blue) gene sets. Genes are ranked on the x-axis from most enriched in cases to most enriched in controls, and position on the y-axis represents significance. The yellow density shows the distribution of 18,000 remaining genes. Our burden signal is driven by a mixture of genes where rare variants are risk increasing (e.g. ***NOD2***) and risk decreasing (*IL23R*).

**Table 1.**
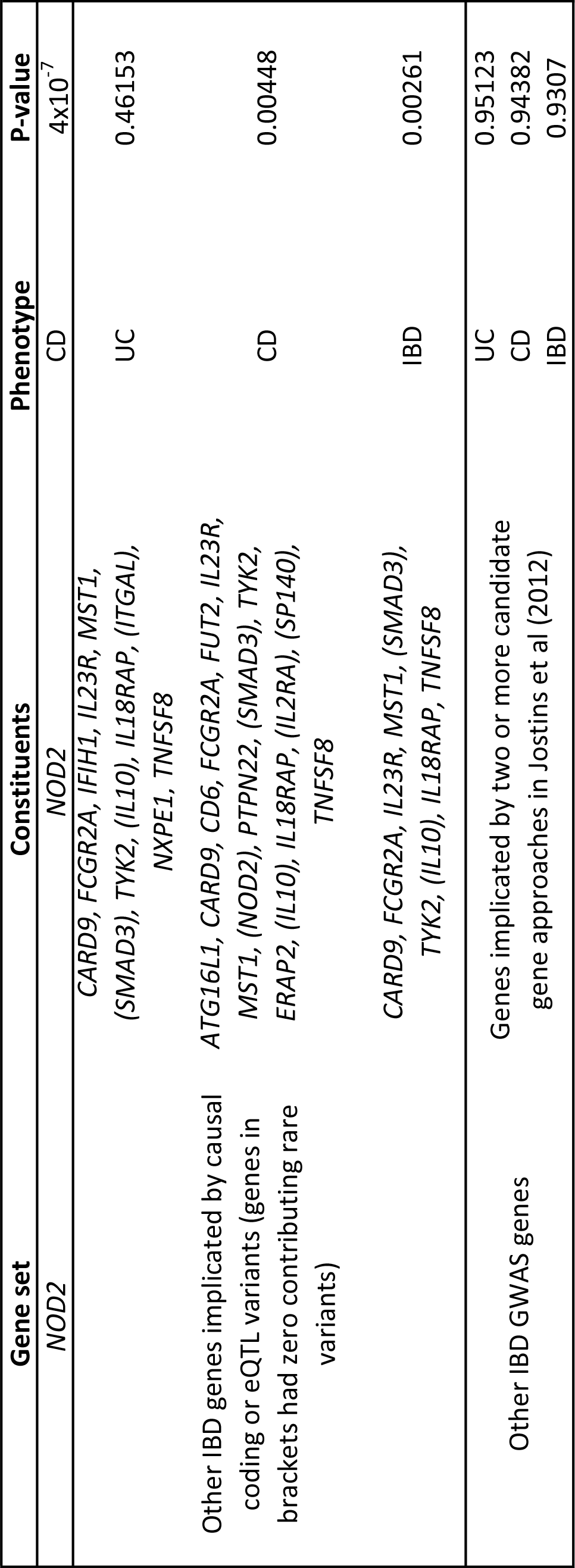
Burden of rare variation in IBD gene sets.

We extended this approach to evaluate rare regulatory variation, using enhancer regions described by the FANTOM5 project. Within each robustly defined enhancer^43^, we tested all observed rare variants, as well as the subset predicted to disrupt or create a transcription factor binding motif^17^. We combined groups of enhancers with cell- and/or tissue-type specific expression, in order to improve power in an analogous fashion to the gene set tests above. However, none of these tissue or cell specific enhancer sets had a significant burden of rare variation after correction for multiple testing (Supplementary Table 13).

## Discussion

We investigated the role of low frequency variants of intermediate effect in IBD risk through a combination of low-coverage whole genome sequencing and imputation into GWAS data. In order to maximize the number of IBD patients we could sequence, and thus our power to detect association, we sequenced our cases at lower depth than the controls available to us via managed access. While joint and careful analysis largely overcomes the bias this introduces, this is just one example of the complexities associated with combining sequencing data from different studies. Such challenges are not just restricted to low coverage whole-genome sequencing designs; variable pulldown technology and sequencing depth in the 60,000 exomes in the Exome Aggregation Consortium^44^ necessitated a simultaneous analysis of such analytical complexity and computational intensity that it would be prohibitive at all but a handful of research centers. Therefore, if rare variant association studies are to be as successful as those for common variants’ computationally efficient methods and accepted standards for combining sequence datasets need to be developed.

We have participated in one such example of a powerful joint analysis by contributing our WGS data to the Haplotype Reference Consortium^26^, which has collected WGS data from more than 32000 individuals into a reference panel that allows accurate imputation down to 0.1% allele frequency. Indeed, one might question the value of generating our own IBD-enriched imputation panel for the analyses described here, when a public resource will eventually offer better accuracy in existing GWAS data. Ultimately, however, most of the data in the Haplotype Reference Consortium comes from projects like ours’ so individual groups must balance investment in sequencing and waiting for development of public resources.

We discovered an association to a low frequency missense variant in ***ADCY7***, which represents the strongest ulcerative colitis risk allele outside of the major histocompatibility complex. The most straightforward mechanistic interpretation of this association is that loss-of-function of ***ADCY7*** reduces production of cAMP, leading to an excessive inflammatory response that predisposes to IBD. Previous evidence suggested that general cAMP-elevating agents that act on multiple adenylate cyclases might, in fact, worsen IBD^45^. While members of the adenylate cyclase family have been considered potential targets in other contexts ^37^, specific upregulation of ***ADCY7*** has not yet been attempted, raising the intriguing possibility that altering cAMP signalling in a leukocyte-specific way might offer therapeutic benefit in IBD.

Despite our study being specifically designed to interrogate both coding and non-coding variation, our sole new association was a missense variant. This is perhaps unsurprising, as the only previously identified IBD risk variants with similar frequencies and odds ratios are protein-altering changes to ***NOD2***, *L23R* and *CARD9*. More generally, the alleles with largest effect sizes at any given frequency tend to be coding^17^, and are therefore the first to be discovered when new technologies expand the frequency spectrum of genetic association studies. This pattern is further reinforced by the tantalizing evidence we found for a burden of very rare coding variants in previously implicated IBD genes. Future deep sequencing studies (exomes or whole genomes), which have near complete power to detect such variants’ have the potential to reveal allelic series ranging from rare highly penetrant variants to common, weak GWAS signals.

Nonetheless, it is likely that nearly all low-frequency IBD susceptibility alleles are regulatory, as is the case for common risk variants, but their effect sizes are too modest to be detected by our current sample size. The paucity of large effects at low frequency variants, modest additional heritability explained by those variants, and the fact that rare variants can hardly ever explain a large fraction of population variation in relatively common diseases, suggest a dichotomy where rare variant association studies are more readily interpreted, but common variant association studies are better suited to discover new biology for a given budget. The ***ADCY7*** association offers a direct window on a new IBD mechanism, but is a relatively meager return compared to the number of loci discovered more simply by increasing GWAS sample size^31^. Together, our discoveries highlight a number of lessons of more general relevance beyond IBD genetics and underline the fact that, while there is much to be learned, transitioning from GWAS to WGS-based studies will not be straightforward.

## Acknowledgements

We would like to thank all individuals who contributed samples to the study. This work was co-funded by the Wellcome Trust [098051] and the Medical Research Council, UK [MR/J00314X/1]. Case collections were supported by Crohn’s and Colitis UK. KMdL, LM, YL, CAL, CAA and JCB are supported by the Wellcome Trust [098051; 093885/Z/10/Z]. KMdL is supported by a Woolf Fisher Trust scholarship. CAL is a clinical lecturer funded by the NIHR. We acknowledge support from the Department of Health via the NIHR comprehensive Biomedical Research Centre awards to Guy’s and St Thomas’ NHS Foundation Trust in partnership with King’s College London and to Addenbrooke’s Hospital, Cambridge in partnership with the University of Cambridge. This research was also supported by the NIHR Newcastle Biomedical Research Centre. The UK Household Longitudinal Study is led by the Institute for Social and Economic Research at the University of Essex and funded by the Economic and Social Research Council. The survey was conducted by NatCen and the genome-wide scan data were analysed and deposited by the Wellcome Trust Sanger Institute. Information on how to access the data can be found on the Understanding Society website https://www.understandingsociety.ac.uk/.

## Competing financial interests

The authors declare no competing financial interests.

## Author contributions

YL, KMdL, LJ, LM, JCB and CAA performed statistical analysis. YL, KMdL, LJ, LM, JCL, CAL, EGS, JR, MaP, SN, and SMC processed the data. TA, CE, NAK, AH, CH, JCM, JCL, CM, WGN, JS, AS, MT, HU, DCW, NJP, CWL, CGW, MP, and CGM contributed samples/materials. YL, KMdL, LM, JCL, MP, CAL, NAK, JCB and CAA wrote the paper. All authors read and approved the final version of the manuscript. JCM, MP, CWL, TA, NJP, JCB and CAA conceived & designed experiments. JCB and CAA jointly supervised the research. YL and KMdL contributed equally to this work.

## Methods

### Preparation of genome-wide genetic data

*Sample ascertainment and sequencing*. 4,686 British IBD cases, diagnosed using accepted endoscopic, histopathological and radiological criteria, were sequenced to low depth (2-4x) using Illumina HiSeq paired-end sequencing. 3,910 population controls, also sequenced to low depth (7x) using the same protocol, were obtained from the UK10K project. Case sequence data was aligned to the human reference used in Phase II of the 1000 Genomes project^46^. Control data was aligned to an earlier human reference (1000 Genomes Phase I)^47^, and then updated to the same reference as the cases using BridgeBuilder, a tool we developed (Supplementary Methods).

*Genotype calling and quality control*. Variants were joint called across 8,424 samples, using samtools and bcftools for SNVs and INDELs, and GenomeSTRiP for structural variants. Structural variants were filtered using standard GenomeSTRiP quality metrics as described in the Supplementary Methods. SNVs were filtered using support vector machines (SVMs) trained on variant quality statistics output from samtools. Each variant was required to pass with a minimum score of 0.01 from at least two out of five independent SVM models. Indels were filtered using GATK VQSR, with a truth sensitivity threshold of 97% (VQSLOD score of 1.0659).

*Genotype refinement and further quality control*. Following initial SNV and INDEL quality control, genotypes at all passing sites were refined via BEAGLE^24^. Variants were then filtered again to remove those showing significant evidence of deviation from Hardy-Weinberg equilibrium (HWE) in controls (P_HWE_<1×10^−7^), a significant frequency difference (P < 1×10^−3^) in samples sequenced at the Wellcome Trust Sanger Institute versus the Beijing Genomics Institute, >10% missing genotypes following refinement (posterior probability < 0.9), SNPs within three base pairs of an INDEL, and allow only one INDEL to pass when clusters of INDELs were separated by two or fewer base pairs. Following these exclusions, a second round of genotype refinement was performed. Sample quality control was then applied to remove samples with an excessive heterozygosity rate (μ ± 3.5*σ*), duplicated or related individuals, and individuals of non-European ancestry (Supplementary Methods).

*Novel GWAS samples*. A further 11,768 British IBD cases and 10,484 population control samples were genotyped on the Human Core Exome v12 chip. Detailed information on ascertainment, genotyping and quality control are described elsewhere^31^.

*Existing GWAS cohorts*. 1748 Crohn’s disease cases and 2936 population controls genotyped on the Affymetrix 500K chip, together with 2361 ulcerative colitis cases and 5417 population controls genotyped on the Affymetrix 6.0 array, were obtained from the Wellcome Trust Case Control Consortium (WTCCC)^27,28^. Both datasets were converted to build 37 using liftOver^48^.

*Imputation*. The whole genome sequences described above were combined with 2504 samples from the Phase 3 v5 release of the 1000 Genomes project (2013-05-02 sequence freeze) to create a phased imputation reference panel enriched in IBD-associated variants. We used PBWT^49^ to impute from this reference panel (114.2 million total variants) into the three GWAS panels described above, after removing overlapping samples. This results in imputed whole genome sequences for 11,987 cases and 15,191 controls.

### Common and low-frequency variation association testing

*Association testing and meta-analysis*. We tested for association to ulcerative colitis, Crohn’s disease and IBD separately within the sequenced samples and three imputed GWAS panels using SNPTEST v2.5, performing an additive frequentist association test conditioned on the first ten principal components for each cohort (calculated after exclusion of the MHC region). We filtered out variants with MAF < 0.1%, INFO < 0.4, or strong evidence for deviations from HWE in controls (p_HWE_<1×10^−7^), and then used METAL (release 2011-03-05) to perform a standard error weighted meta-analysis of all four cohorts. Only sites for which all cohorts passed our quality control filters were included in our meta-analysis.

*Quality control*. The output of the fixed-effects meta-analysis was further filtered, and sites with high evidence for heterogeneity (I^2^>0.90) were discarded. In addition, we discarded all genome-wide significant variants for which the meta-analysis p-value was not lower than all of the cohort-specific p-values. Finally, and in order to minimise the false positive associations due to mis-imputation, sites which did not have an info score ≥ 0.8 in at least three of the four datasets (two of the three for Crohn’s disease and ulcerative colitis) were removed.

*Locus definition*. A linkage disequilibrium (LD) window was calculated for every genome-wide significant variant in any of the three traits (Crohn’s disease, ulcerative colitis, IBD), defined by the left-most and right-most variants that are correlated with the main variant with an r^2^ of 0.6 or more. The LD was calculated in the GBR and CEU samples from the 1000 Genomes Phase 3, release v5 (based on 20130502 sequence freeze and alignments). Loci with overlapping LD windows, as well as loci whose lead variants were separated by 500kb or less, were subsequently merged, and the variant with the strongest evidence of being associated was kept as the lead variant for each merged locus. This process was conducted separately for each trait. A locus was annotated as known when there was at least one variant in it that was previously reported to be of genome-wide significance (irrespective of the LD between that variant and the most associated variants in the locus). Otherwise, a locus was annotated as putatively novel.

*Conditional analysis*. Conditional analyses were conducted using SNPTEST 2.5, as for the single variant association analysis. P-values were derived using the score test (default in SNPTEST v2.5). In order to fully capture the ***NOD2*** signal when investigating the remaining signal in the region, we conditioned on seven variants which are known to be associated: rs2066844, rs2066845, rs2066847, rs72796367, rs2357623, rs184788345, and rs104895444.

*Heritability explained*. The SNP heritability analysis was performed on the dichotomous case-control phenotype using constrained REML in GCTA39 with a prevalence of 0.005 and 0.0025 for Crohn’s disease and ulcerative colitis respectively. Hence, all reported values of h2g are on the underlying liability scale. To further eliminate spurious associations, we computed genetic relationship matrices (GRMs) restricted to all variants with MAF ≥ 0.1%, imputation r^2^ ≤ 0.6, missing rate ≤ 1% and Hardy-Weinberg equilibrium P-value ≥ 1×10^−7^ in controls for each GWAS cohort. We further checked the reliability and robustness of our estimates by performing a joint analysis across all autosomes, a joint analysis between common (MAF≥1%) and rare variants (0.1%≥MAF<1%), and LD-adjusted analysis using LDAK^50^ (Supplementary Methods).

### Rare variation association testing

*Additional variant quality control*. Additional site filtering was used, as rare sites are more susceptible to differences in read depth between cases and controls (Supplementary Figure 8). This included removing singletons, as well as sites with: missingness rate > 0.9 when calculated using genotype probabilities estimated from the samtools genotype quality (GQ) field; low confidence observations comprising ≥ 1% of non-missing data, or; INFO < 0.6 in the appropriate cohorts.

*Association testing*. Individual gene and enhancer burden tests were performed using an extension of the Robust Variance Score statistic^42^ (Supplementary Methods), to adjust for the systematic coverage bias between cases and controls. This required the estimation of genotype probabilities directly from samtools (using the genotype quality score), as genotype refinement using imputation results in poorly calibrated probabilities at rare sites. Burden tests were performed across sites with a MAF ≤ 0.5% in controls and within genes defined by Ensembl, or enhancers as based on its inclusion in the FANTOM5 ‘robustly-defined, enhancer set^43^. For each gene, two sets of burden tests were performed: all functional coding variants and all predicted damaging (CADD > 21) functional coding variants (Supplementary Table 6). For each enhancer, burden tests were repeated to include all variants falling within the region, and just the subset predicted to disrupt or create a transcription factor binding motif (Supplementary Methods).

*NOD2 independence testing*. We evaluated the independence of the rare NOD2 signal from the known common coding variants in this gene (rs2066844, rs2066845, and rs2066847). Individuals with a minor allele at any of these sites were assigned to one group, and those with reference genotypes to another. Burden testing was performed for this new phenotype in both variant sets that contained a significant signal in Crohn’s disease vs controls.

*Set definition*. The individual burden test statistic was extended to test across sets of genes and enhancers using an approach based on the SMP method^41^, whereby the test statistic for a given set is evaluated against the statistics from the complete set (e.g. all genes), to account for residual case-control coverage bias. The sets of genes confidently associated with IBD risk were defined based on implication of specific genes in ulcerative colitis’ Crohn’s disease or IBD risk through fine-mapping, eQTL and targeted sequencing studies (Supplementary Table 9). The broader set of IBD genes was defined as any remaining genes implicated by two or more candidate gene approaches in Jostins et al (2012)^51^. Enhancer sets were defined as those showing positive differential expression in each of 69 cell types and 41 tissues’ according to Andersson et al^43^ (Supplementary Table 11).

